# *In silico* study for decoding the correlated role of *MCM7* gene in progression of breast cancer and Alzheimer’s disorder

**DOI:** 10.1101/2021.06.13.448221

**Authors:** Navneeth Sriram, Sunny Mukherjee, Mahesh Kumar Sah

**Affiliations:** Department of Biotechnology, Dr. B. R. Ambedkar National Institute of Technology, Jalandhar, Punjab-144011, India

**Keywords:** Breast cancer, Alzheimer’s disorder, CD209, MCM7, gene annotation, pathway analysis

## Abstract

Breast cancer and Alzheimer’s disease (AD) are two of the progressive and detrimental disorders affecting large population around the globe. While the chemotherapy of breast cancer is well established and enriched, the AD still lacks it due to unrecognized peripheral biomarkers for detection and targeted therapy. This study aimed to identify common molecular signature markers in breast cancer (grade 1, 2, and 3) and AD for the diagnosis and prognosis. We used two microarray datasets (GSE42568, GSE33000) respectively for both disorders that led to identification of two common differentially expressed genes (DEGs), namely MCM7 and CD209, as common players in both these two conditions. While the pattern of expression of CD209 gene running upregulated in both disorders, the MCM7 showed unusual contrary in its pattern of expression. The expression of MCM7 is downregulated in breast cancer but upregulated in AD. Gene set and protein overrepresentation analysis, protein-protein interaction (PPI), and protein subcellular localizations analyses of this underrated MCM7 gene was performed with further prediction and validation of its structure. The findings may pave the way in designing therapeutic approaches to ameliorate AD.

## 1. INTRODUCTION

The modern technologies employed for delivering research and clinical treatment for cancer and neurodegenerative disorders have brought the diagnosis, prognosis, and treatment more effective and personalized. While earlier, those were based on clinical-pathological observations, the present in silico assisted research revolutionized the efficacy and discovery of diagnostic and therapeutic agents. Among wide range of cancer types, breast cancer dominant *w.r.t*. high mortality and growth rate of new women patients. According to WHO report, in 2020, there were 2.3 million women diagnosed with breast cancer and 685 000 deaths globally. The subtypes of breast cancer such as ductal and lobular carcinoma, oestrogen receptor-positive (luminal A), HER2^+^ (Human Epidermal Growth Factor Receptor 2), triple negative and basal-like are common [1]. The histological grading of breast cancer indicates the extent of malignancy in a patient are grade 1 (well differentiated cancer cells growing slowly), grade 2 (moderately differentiated cancer cells) and grade 3 (poorly differentiated cancer cells growing faster)[2]. An exhaustive cancer research encompasses the use of microarray and sequencing-based technologies to analyse gene expression, to evaluate pathogenesis, and predict prognostic factors for various tumors[3]. The importance of the DNA replication licensing factors or MCMs in promoting expression coupled with altered phosphorylation of p53 that leads to drug resistance is reported earlier[4]. It is reported that at a cellular level, there is an inverted dynamic that happens between Cancer and Alzheimer’s disease (AD), a complex and neurodegenerative disorder. AD is a common form of dementia in elders (>65 years), leading to cognitive impairment and memory loss and occurs due to neuron cells death, atrophy, and/or inflammation in hippocampus region of brain [5]. At a molecular level, the condition is primarily known by detecting the accumulation of amyloid β plaques, NFTs (Neurofibrillary Tangles) and the neurites containing the hyperphosphorylated tau protein [6]. The most common mutation that was found in them was the mutation in ε4 allele of the ApoE gene and it increased the genetic risk factor for sporadic AD by 95% [7]. The most common genes whose mutation greatly increases the susceptibility to AD are APP (Amyloid Precursor protein), PS-1 (Presenilin-1) and PS-2 (Presenilin-2) respectively [8]. In addition, genome-wide association studies (GWASs) have identified several genes as potential risk factors for AD, such as *clusterin* (*CLU*), *complement receptor 1* (*CR1*), *phosphatidylinositol binding clathrin assembly protein* (*PICALM*), and *sortilin-related recepto*r (*SORL1*) [9]. The recent inclusions are novel genes that might be engaged in triggering late-onset D, such as *triggering receptor expressed on myeloid cells 2* (*TREM2*) and *cluster of differentiation 33* (*CD33*) [9]. The identification of new AD-related genes plays vital role in unravelling the pathogenesis mechanisms that may guide the discovery of new treatment strategies.

As reported earlier, in Cancer, there is a state of sustained cell signalling that results in its proliferation, whereas in AD, the proliferative signalling is impaired [10]. The cancer cells tend to evade the growth suppressors but these have a very strong effect on the AD-afflicted neurons [10]. The apoptotic processes are downregulated in Cancer while they are upregulated in AD [11]. Due to this property, studies are being carried out on the repurposing of anti-cancer drugs in order to cover both the conditions [12]. This is further supported by the finding that the patients who underwent chemotherapy had a lower risk of AD than those who hadn’t undergone chemotherapy [13]. In the present work, the common genes for both breast cancer and the AD is analysed and their pattern of expression are further studied for possible repurposing of anti-cancer drugs of breast cancer for the treatment of AD. The article focusses on the effect of both the conditions on decoded gene functioning and to establish the regulatory effect of associated protein on AD.

## 2. EXPERIMENTAL DESIGN

The datasets for Cancer and Alzheimer’s Disease were collected from GEO database of NCBI (https://www.ncbi.nlm.nih.gov/geo/) [14]. These datasets were compared on the basis of normal controls versus diseased conditions. To statistically analyze the data, the data were normalized and log transformation was applied. Further, Benjamini-Hochberg correction was used to eliminate the false discovery rate. The obtained output and the list of significant differentially expressed genes (DEGs) were downloaded (P_adj_<0.01). The genes in common were then identified with the help of VENNY 2.1.0, a Venn Diagram generating tool (https://bioinfogp.cnb.csic.es/tools/VENNY2.1.0/). The gene ontology and gene enrichment studies were conducted using Gene Ontology Resource (http://geneontology.org/) [15] and ENRICHR [16] respectively. Further, the analysis of protein-protein interaction of the target protein and their respective sub-cellular localization were determined. The domain analysis of the target gene, prediction of its structure followed by the validation of the analysis were performed. The workflow for the present methodology is depicted in ***Figure 1***.

**Figure 1:**
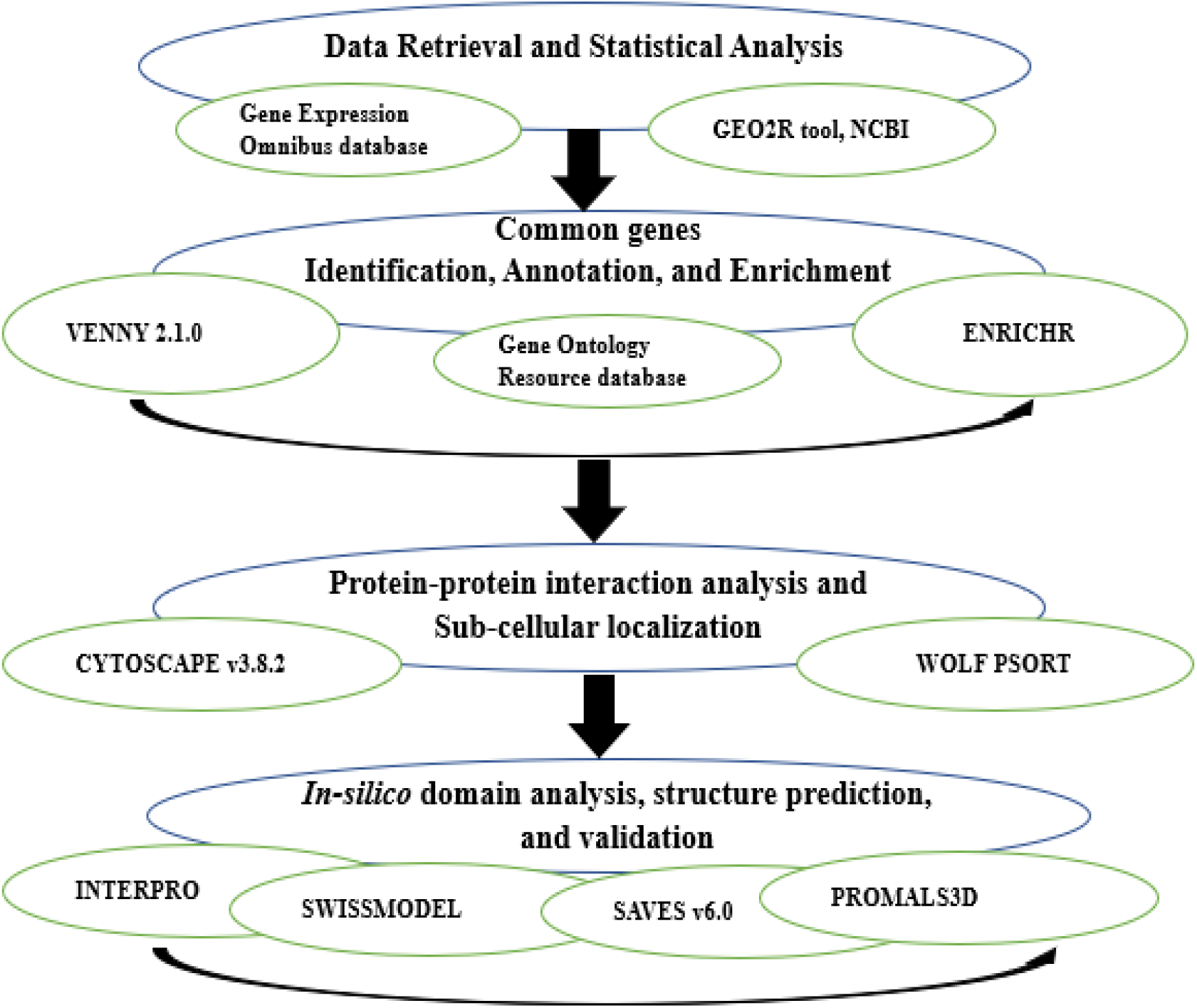
The flowsheet of methodology for the identification of potential genes associated with breast cancer and Alzheimer’s disorder in common.

### 2.1. Datasets and identification of DEGs from microarray datasets

For collecting the DEGs associated with Breast Cancer, the gene expression profiles were studied (GSE42568) [17]. The total number of samples were 121, out of which 17 were controls, 11 were of Grade 1, 40 were of Grade 2 and 53 were of Grade 3. For collecting the DEGs associated with AD, the gene expression profile of human prefrontal cortex brain tissues (GSE33000) was analysed [18]. The total number of present samples were 621, out of which 467 from the AD patients while the rest 154 were samples from HD (Huntington’s Disease) patients. Among 467, 135 were of Male AD, 123 were Male control, 175 were Female AD, and 34 were Female controls. Those 467 samples were analyzed for the study.

These samples were first selected, assigned a group, and then GEO2R tool was used to analyze the data. Benjamini-Hochberg correction was applied to the analysed data to eliminate the false discovery and further the data was normalized by log transformation. To further analyze the data, the LIMMA precision weights (Linear Models for Microarray Analysis) was also applied (P<0.01). After applying all these modifications, the data was analyzed. To obtain the DEGs for these dataset, the plot displays were also selected, which were Breast cancer control vs Breast Cancer Grade 1; Breast Cancer Control vs Breast Cancer Grade 2 and Breast Cancer Control vs Breast Cancer Grade 3, Female AD vs Female Control and Male AD vs Male Control, respectively.

### 2.2. Gene ontology and enrichment study

In order to conduct gene ontology study, Gene Ontology Resource (http://geneontology.org/) web tool was used. The genes that were common in both the datasets were firstly identified with VENNY 2.1.0 (https://bioinfogp.cnb.csic.es/tools/VENNY2.1.0/), a Venn Diagram generating tool. After identification, the common genes were searched and analysed for enrichment using ENRICHR online tool.

### 2.3. Protein-protein interaction analysis

The Protein-Protein interaction (PPI) network of the target protein unveiled after enrichment analysis, MCM7, was analysed using Cytoscape v3.8.2 [19]. The STRING plug-in of Cytoscape v3.8.2 was used to perform an automatic search for known interacting partners of target protein under study using data from the STRING network database [20]. Search parameters were set as follows: confidence score set at highest (0.99), additional interactors set at maximum (100) and smart delimiters were activated. Nodes represent the proteins and the edges represent the interactions among them. After the network was imported into Cytoscape, the individual interactions of each nodes were observed and image data was exported for further analyses.

### 2.5 Determination of the subcellular localization of interacting proteins

The WoLF PSORT [21] online server in combination with experimentally validated data from UniProt [22] were used to predict the subcellular localization of all the proteins present in the interaction network of target protein. The prediction of the subcellular localization is useful for determination of protein functions as well as the feasibility of a protein involved in the interaction network (direct/via cascade) to serve as a potential target during the drug discovery process.

### 2.6 *In-silico* domain analysis and structure prediction of target protein

Domain analysis is necessary for the prediction of active or functional sites on a protein. Since these are the regions that pharmaceutical ligands often bind to, those can be potentially served as drug targets. The domain analysis of target protein was performed using European Bio-Informatics Institute (EBI) INTERPRO [23] server after extraction of the FASTA sequence from UniProt. The secondary structure prediction of a protein, in order to understand its folded state as well as the presence of any loops or turns, was made using UCLCS Bio-Informatics group’s PSIPRED (PSI-BLAST based secondary structure PREDiction) [24] online tool. 3D homology modelling was performed using SWISSMODEL [25] in order to validate pre-existing structure from RCSB PDB [26] and for the observation of various reactive amino acid groups. The predicted model was validated using the SAVES v6.0 utilizing the VERIFY3D [27] and PROCHECK [28]. Finally, all the predicted structures and published structures were compared for sequence similarity using PROMALS3D [29] server, identical sequences would conclusively validate the PDB structure for target protein.

## 3. RESULTS AND DISCUSSION

### 3.1. Identification of DEGs

After conducting the statistical analyses using GEO2R tools of NCBI, the Volcano plots of the four conditions against their respective controls was obtained ***(Figure 2)***. After obtaining the volcano plots, the significant genes were downloaded and analysed using VENNY to find out the common genes that were present in the four conditions. After getting the common genes, the network was designed using CYTOSCAPEv3.8.2 where the networks files were imported from STRING database. A high confidence (>90%) was set as a parameter to obtain the network.

**Figure 2:**
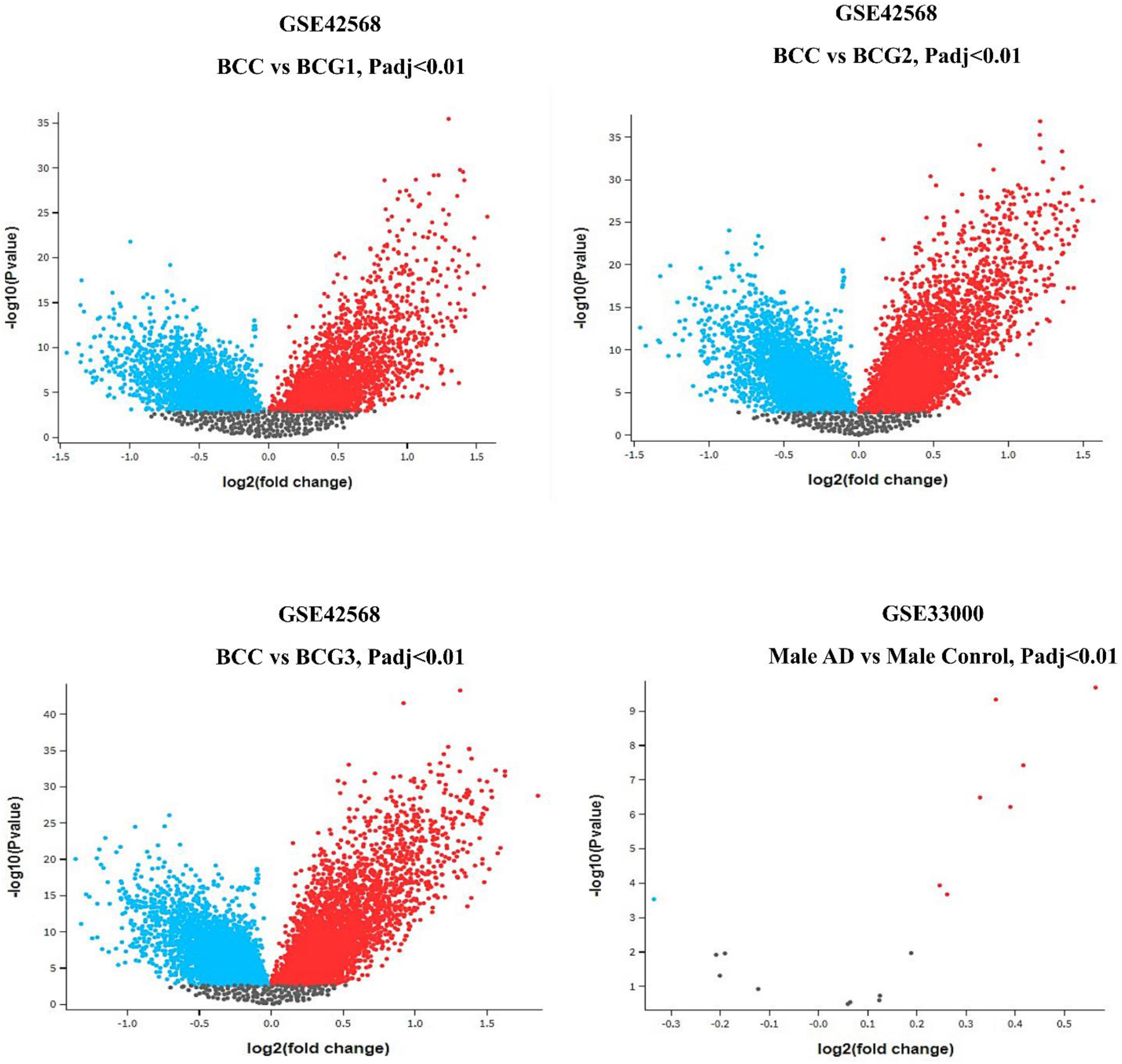
Volcano plots of DEGs between breast cancer Grade 1 (A), Grade 2 (B), Grade 3 (C) and normal breast cancer tissue. Volcano plot of DEGs between Alzheimer’s disordered and normal brain tissues (d). Red dots, significantly upregulated genes in breast cancer tissues; blue dots, significantly downregulated genes in breast cancer tissues; grey dots, not significantly expressed genes. Adj. P< 0.01 was considered to indicate a statistically significant difference.

After analysing the microarray data, it was determined that there were 4091 significant DEGs in Breast Cancer Grade 1, 6548 in Breast Cancer Grade 2 and 7081 in Breast Cancer Grade 3 and in Alzheimer’s Disease dataset, there were 5 DEGs that were significant. Volcano plot was made where x-axis represented log2(fold change) and the y-axis represented the negative log (fold change). The red points denoted the genes whose fold change was greater than 0 and the blue points denoted the genes whose fold change was less than 0. There were no significant genes for Male AD vs Male Control data. Upon determining the DEGs that were common in both Alzheimer’s Disease and Breast Cancer datasets, it was found that two genes, namely, CD209 and MCM7 were the only ones present in both of them with CD209 showing a consistent downregulation pattern across the three grades of breast cancer and Alzheimer’s Disease and MCM7 showing a consistent downregulation pattern against the three grades of breast cancer and an upregulation in case of Alzheimer’s Disease. The gene MCM7 is chosen for further analyses.

Besides, Venny 2.1.0 was used to identify overlapping DEGs across the 4 datasets ***(Figure 3, Table 1)***. Upon analysing the annotations obtained for MCM7, it is found that in case of biological process, it is involved in cell division(GO:0000278) and DNA replication(GO:0006260); in case of molecular function, it is involved in the initial stages of DNA replication (GO:1990518) and in case of cellular compartment, this gene is present as a component of different complexes, most notably being protein-DNA complex(GO:0032993), DNA replication preinitiation complex(GO:0031261) and also is present as a component of nuclear chromosomes (GO:0000228). Upon conducting the enrichment analysis, it is found that the fold enrichment of MCM7 exceeds 100 in cases of initiation of DNA replication (R-HSA-176974) and DNA replication (R-HSA-69306) processes. This also happens in G1/S transitions (R-HSA-69206), G2/M checkpoints (R-HSA-69481) and S phase (R-HSA-69242). The fold enrichment is less than 100 in case of progression of cell cycle (R-HSA-69278), which is consistent with the involvement of MCM7 in replication process.

**Table 1:** Fold change and F-value of common genes

| Gene Symbol | Gene Title | Log2(fold change) |  |  |  | F-value |  |  |  |
| --- | --- | --- | --- | --- | --- | --- | --- | --- | --- |
|  |  | BCG1 | BCG2 | BCG3 | AD | BCG1 | BCG2 | BCG3 | AD |
| MCM7 | Minichromosome Maintenance Complex Component 7 | -0.176 | -0.149 | -0.207 | 0.399 | 3.471 | 3.908 | 7.155 | 6.407 |
| CD209 | CD209 molecule | 0.917 | 0.862 | 0.844 | 0.366 | 16.766 | 23.925 | 24.647 | 9.523 |

**Figure 3:**
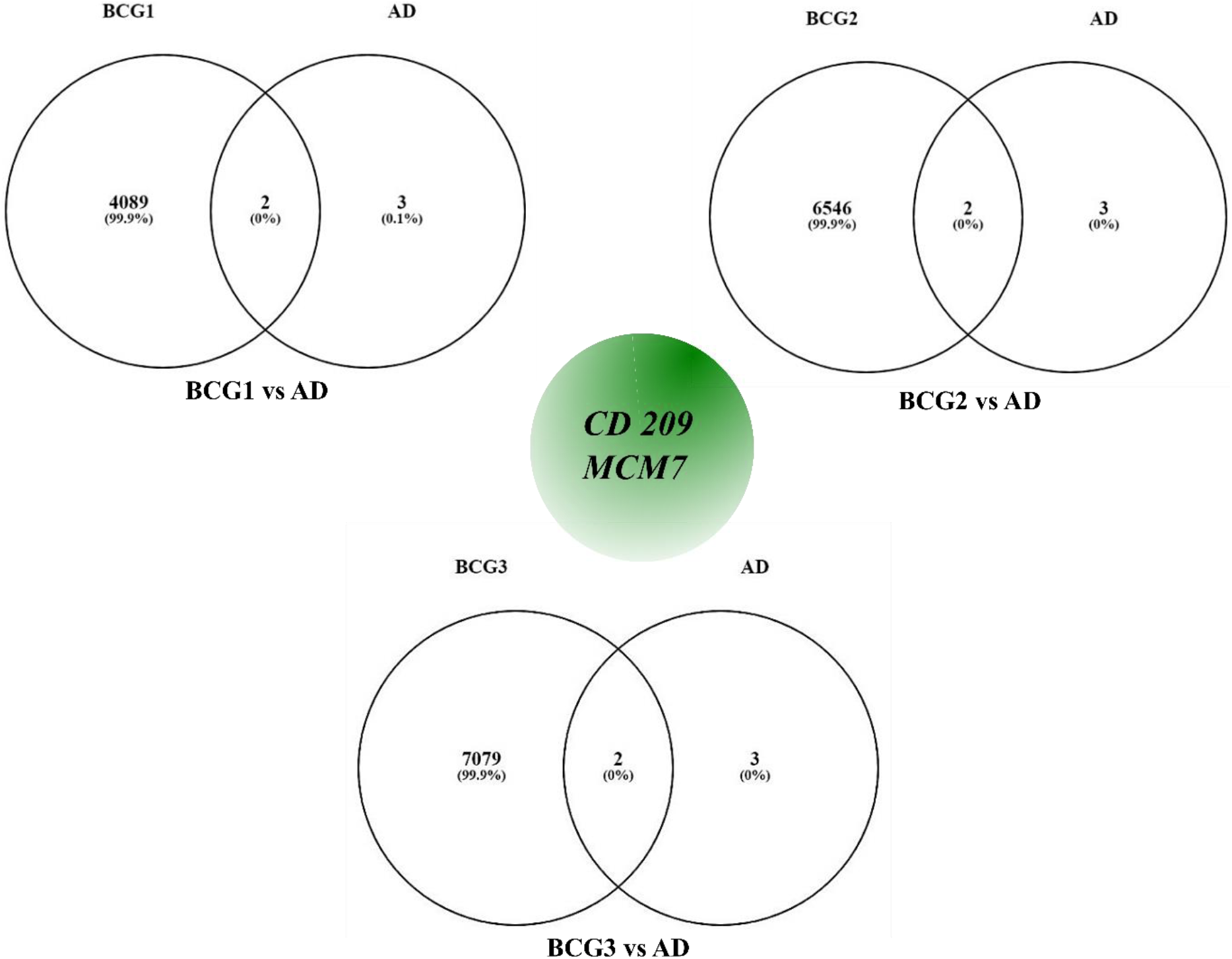
Venn diagram of overlapping DEGs from the GSE42568 (breast cancer grade 1, grade 2, and grade 3) and GSE330000 datasets (Alzheimer’s disorder). DEGs, differentially expressed genes.

The complete PPI network of MCM7 after analysis through Cytoscape revealed 51 nodes and 363 edges. In this case, Nodes represent the proteins and the edges represent the interactions among them. Upon performing a search for the first neighbours (undirected) of our target protein MCM7, it was observed that MCM7 has direct interactions with 35 proteins. It was also observed that 15 other proteins interact with MCM7 using one or the other of the 35 directly interacting protein partners as an intermediate using a cascade mechanism. The cellular function of the interacting partners of MCM7 protein as well as the interaction type (direct/indirect) are elucidated in ***Table 4***.

**Table 2:** Gene Annotation for Biological Process of MCM7 (Gene Ontology Resource)

| Category | GO ID | Term | P-value |
| --- | --- | --- | --- |
| **Biological Process** | GO:1902299 | pre-replicative complex assembly involved in cell cycle DNA replication | 0.000437 |
|  | GO:0006267 | pre-replicative complex assembly involved in nuclear cell cycle DNA replication | 0.000437 |
|  | GO:0000727 | double-strand break repair via break-induced replication | 0.000631 |
|  | GO:0006268 | DNA unwinding involved in DNA replication | 0.000825 |
|  | GO:0006271 | DNA strand elongation involved in DNA replication | 0.000971 |
|  | GO:0022616 | DNA strand elongation | 0.00121 |
|  | GO:0006270 | DNA replication initiation | 0.00214 |
|  | GO:0071364 | cellular response to epidermal growth factor stimulus | 0.00223 |
|  | GO:0070849 | response to epidermal growth factor | 0.00243 |
|  | GO:0033260 | nuclear DNA replication | 0.00286 |
|  | GO:0044786 | cell cycle DNA replication | 0.00291 |
|  | GO:0036388 | pre-replicative complex assembly | 0.0032 |
|  | GO:0000724 | double-strand break repair via homologous recombination | 0.00495 |
|  | GO:0000725 | recombinational repair | 0.00505 |
|  | GO:0032508 | DNA duplex unwinding | 0.0052 |
|  | GO:0032392 | DNA geometric change | 0.00554 |
|  | GO:0000082 | G1/S transition of mitotic cell cycle | 0.00612 |
|  | GO:0044843 | cell cycle G1/S phase transition | 0.00621 |
|  | GO:0071466 | cellular response to xenobiotic stimulus | 0.00631 |
|  | GO:0009410 | response to xenobiotic stimulus | 0.00665 |
|  | GO:0006261 | DNA-dependent DNA replication | 0.00947 |
|  | GO:0006302 | double-strand break repair | 0.00986 |
|  | GO:0006310 | DNA recombination | 0.0109 |
|  | GO:0065004 | protein-DNA complex assembly | 0.0124 |
|  | GO:0006260 | DNA replication | 0.0132 |
|  | GO:0044772 | mitotic cell cycle phase transition | 0.0139 |
|  | GO:0044770 | cell cycle phase transition | 0.0143 |
|  | GO:0071824 | protein-DNA complex subunit organization | 0.0143 |
|  | GO:0071103 | DNA conformation change | 0.0152 |
|  | GO:0042493 | response to drug | 0.0186 |
|  | GO:0008283 | cell population proliferation | 0.0232 |
|  | GO:0071363 | cellular response to growth factor stimulus | 0.0245 |
|  | GO:0006281 | DNA repair | 0.0258 |
|  | GO:0070848 | response to growth factor | 0.0259 |
|  | GO:1903047 | mitotic cell cycle process | 0.0333 |
|  | GO:0000278 | mitotic cell cycle | 0.037 |
|  | GO:0006259 | DNA metabolic process | 0.038 |
|  | GO:0006974 | cellular response to DNA damage stimulus | 0.0383 |
|  | GO:0034622 | cellular protein-containing complex assembly | 0.0418 |
| **Molecular**<br>**Function** | GO:1990518 | single-stranded 3'-5' DNA helicase activity | 0.000243 |
|  | GO:0043138 | 3'-5' DNA helicase activity | 0.000874 |
|  | GO:0017116 | single-stranded DNA helicase activity | 0.000971 |
|  | GO:0003688 | DNA replication origin binding | 0.00117 |
|  | GO:0003678 | DNA helicase activity | 0.00369 |
|  | GO:0003697 | single-stranded DNA binding | 0.00578 |
|  | GO:0004386 | helicase activity | 0.00777 |
|  | GO:0140097 | catalytic activity, acting on DNA | 0.0103 |
|  | GO:0016887 | ATPase activity | 0.0237 |
|  | GO:0017111 | nucleoside-triphosphatase activity | 0.042 |
|  | GO:0016462 | pyrophosphatase activity | 0.0447 |
|  | GO:0016818 | hydrolase activity, acting on acid anhydrides, in phosphorus-containing anhydrides | 0.0448 |
|  | GO:0016817 | hydrolase activity, acting on acid anhydrides | 0.0448 |
| **Cellular**<br>**Component** | GO:0071162 | CMG complex | 0.000534 |
|  | GO:0031261 | DNA replication preinitiation complex | 0.000631 |
|  | GO:0042555 | MCM complex | 0.000631 |
|  | GO:0000781 | chromosome, telomeric region | 0.00728 |
|  | GO:0032993 | protein-DNA complex | 0.00869 |
|  | GO:0000228 | nuclear chromosome | 0.0111 |
|  | GO:0098687 | chromosomal region | 0.0163 |

**Table 3:**
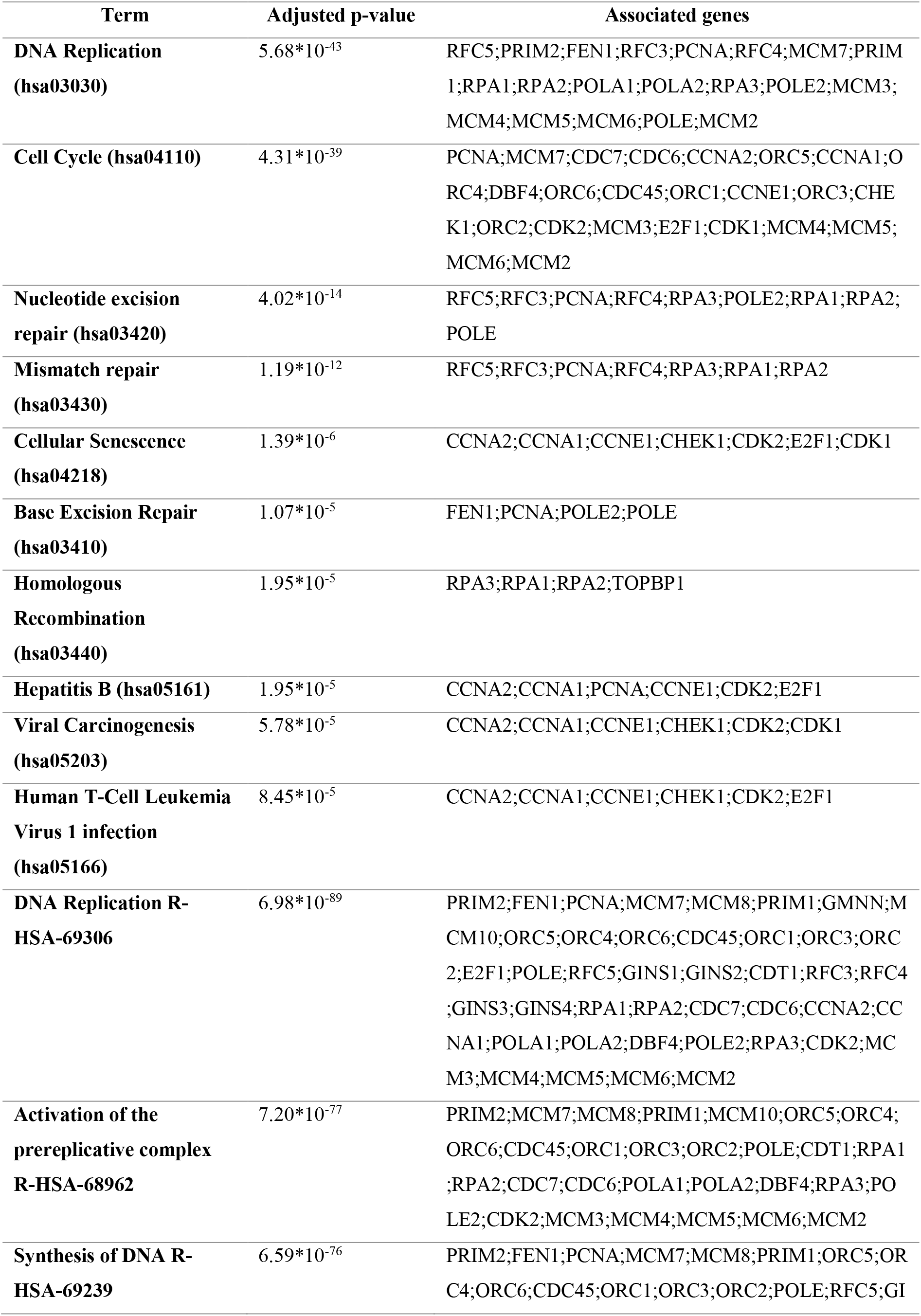
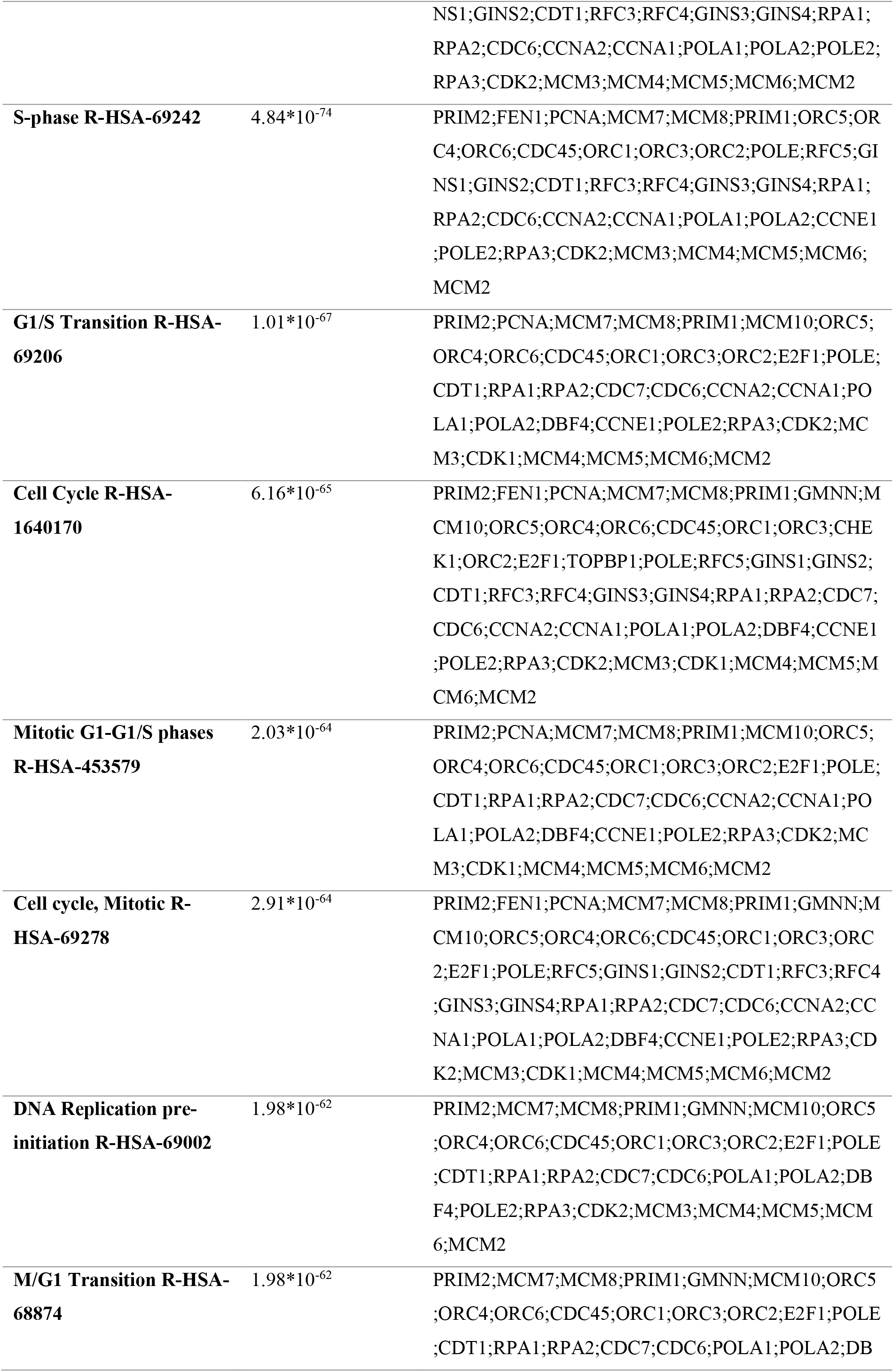
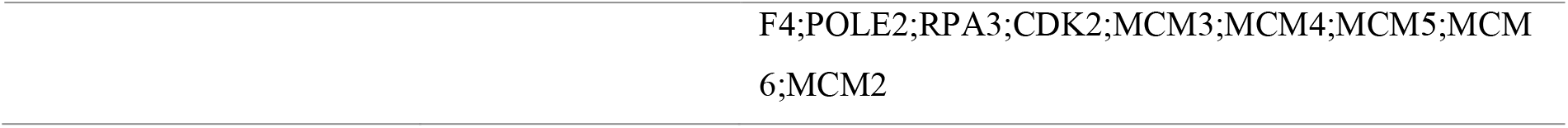
Gene Enrichment Analysis of MCM7

**Table 4:**
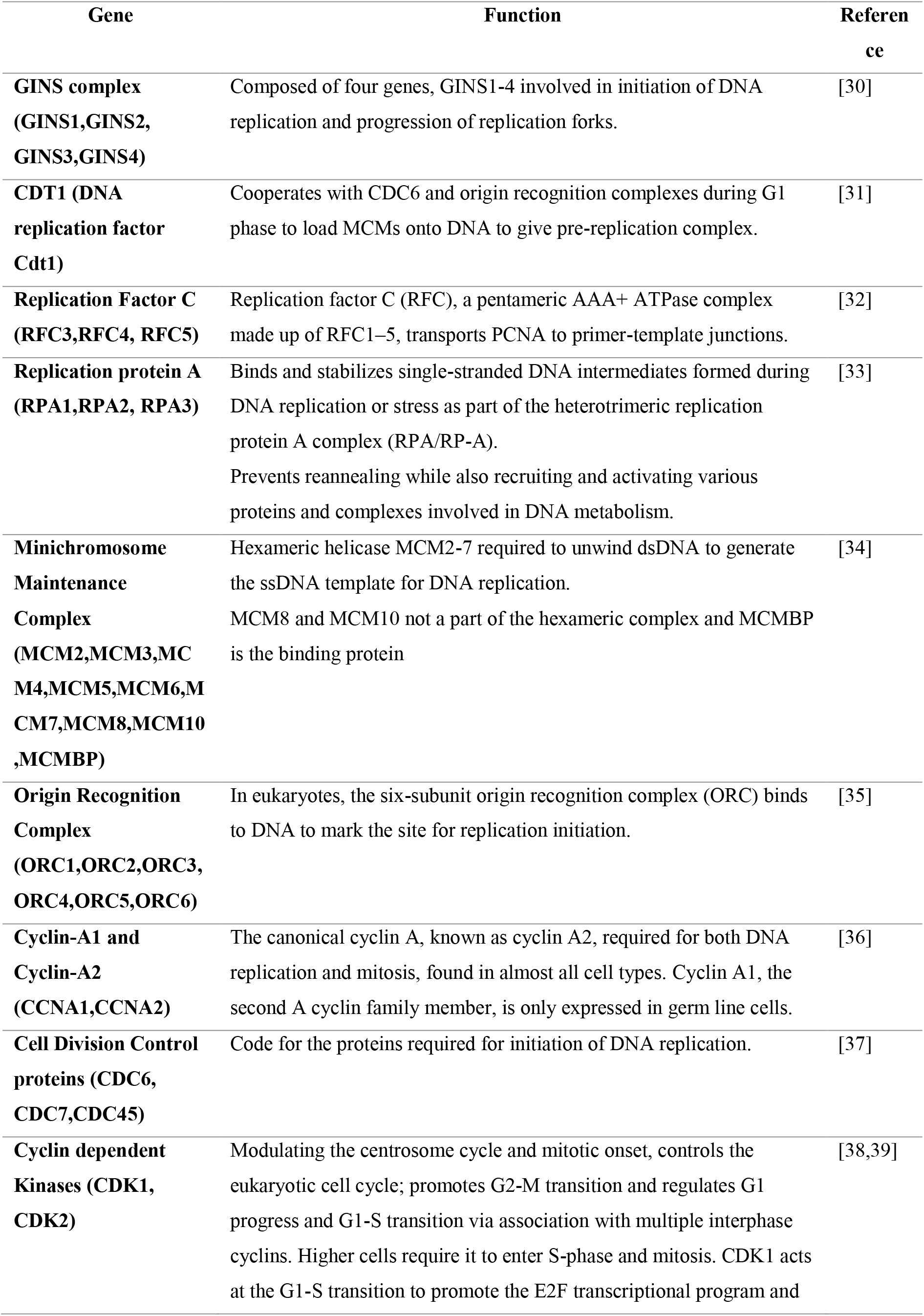
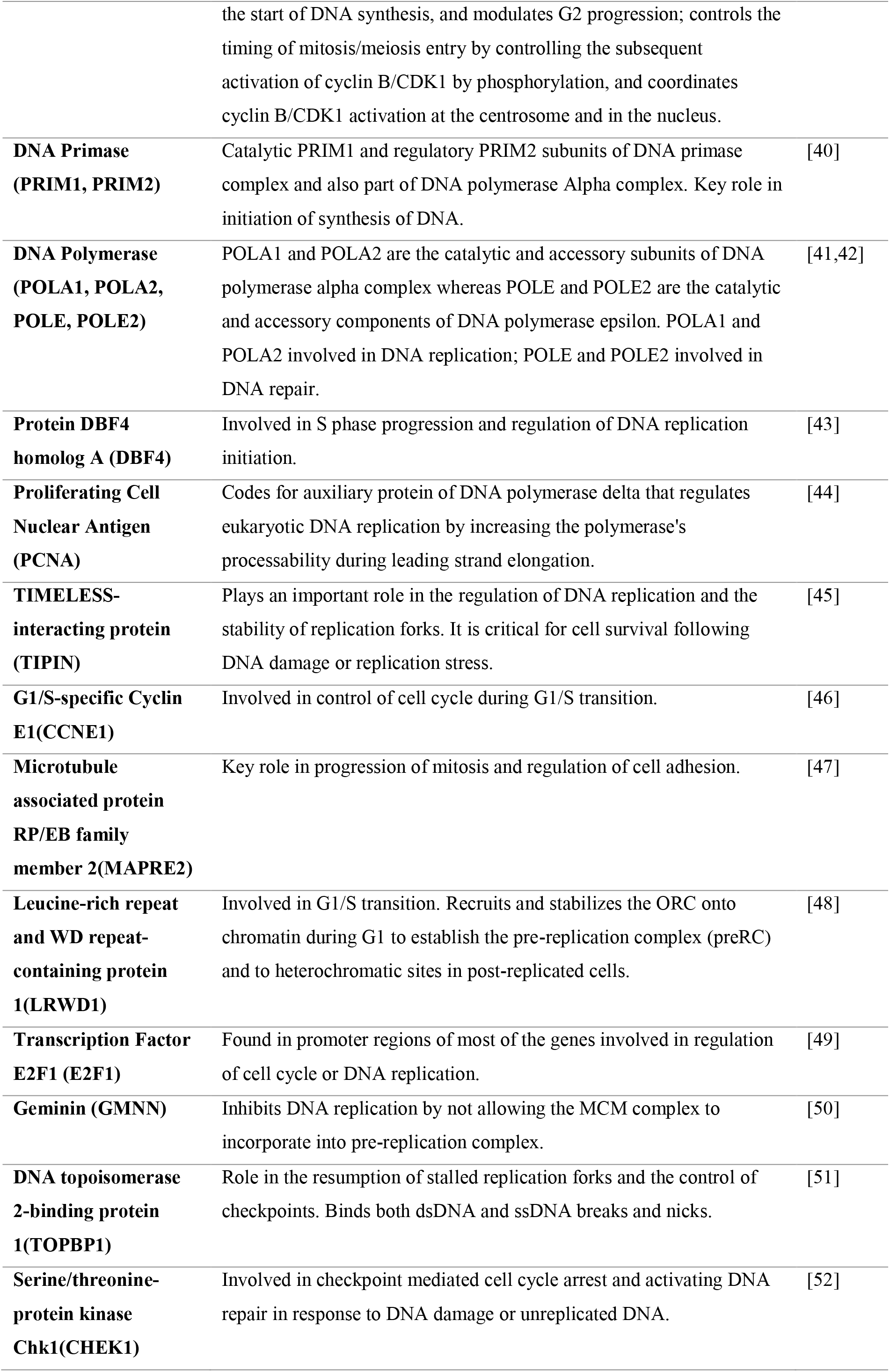
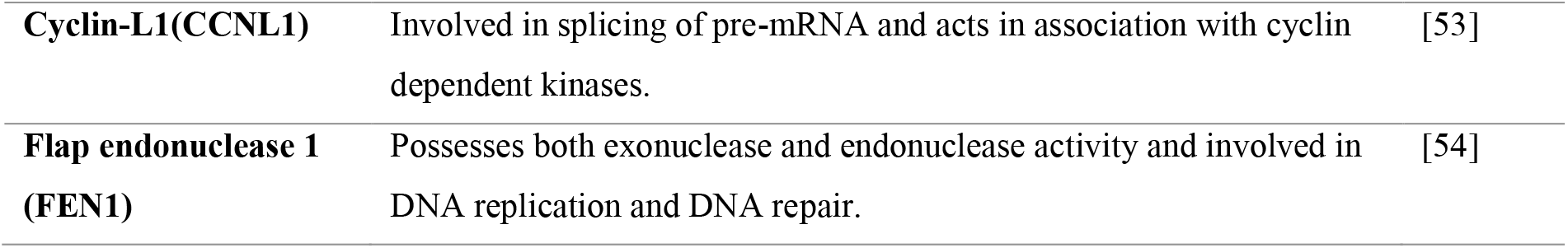
Description of cellular functions of all the proteins associated with the PPI network of MCM7

***Figure 4*** represents the Protein-protein interaction (PPI) network of MCM7 obtained using Cytoscape v3.8.2 (Figure 4A). Nodes represent the proteins and the edges represent the interactions among them. Yellow octagonal node illustrates the protein of interest which has been selected as the central hub. Red octagonal nodes portray the first neighbours (undirected) of MCM7 protein. Blue elliptical nodes depict those proteins that are interacting with MCM7 via another protein using a cascade mechanism. Figure 4B displays the PPI Network between MCM7 and its first neighbours (undirected) (isolated from Fig 4(A) network). Highest confidence score (0.99), maximum additional interactors (100) and smart delimiters were the parameters used for the construction of this PPI network.

**Figure 4:**
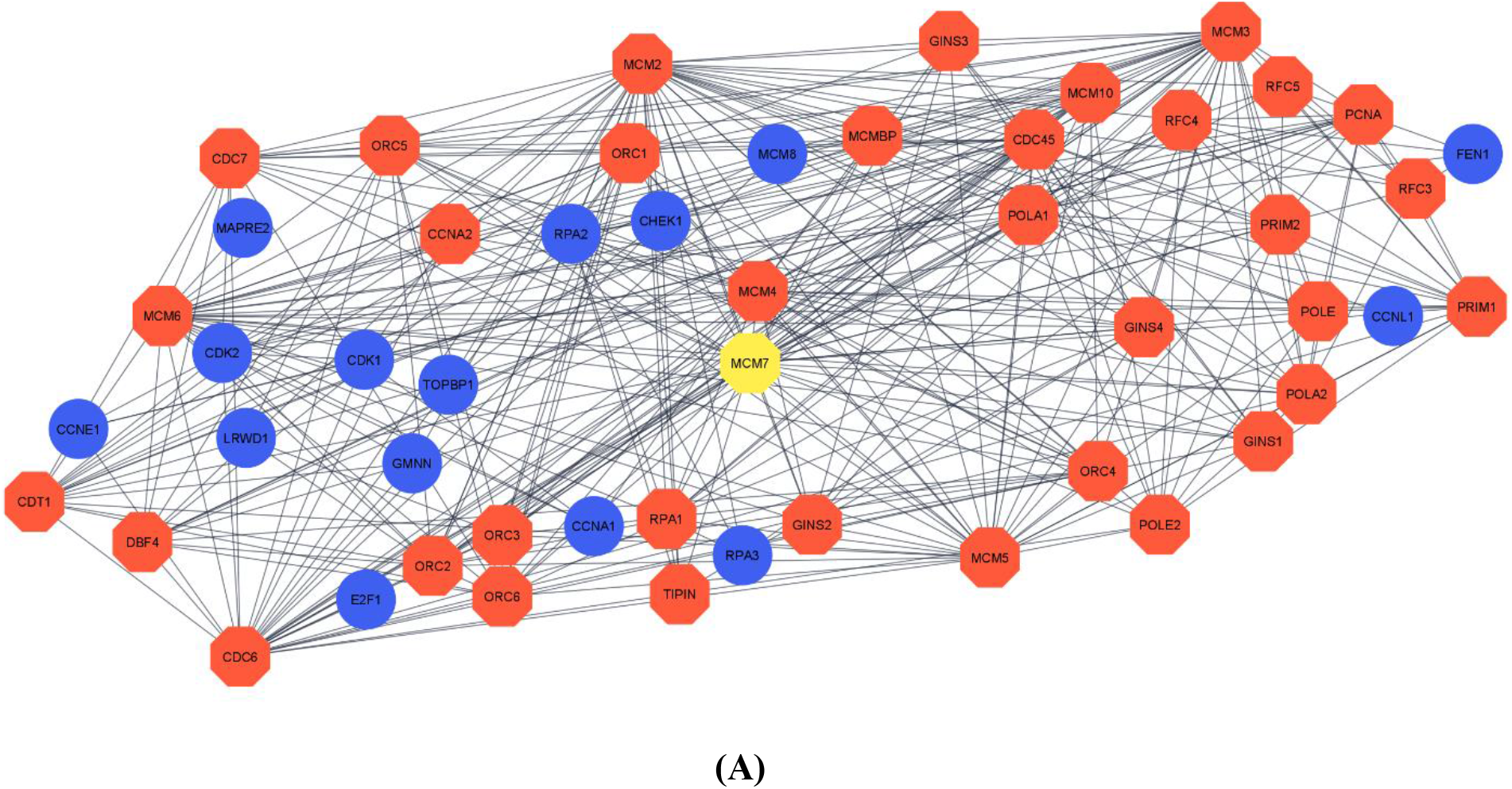
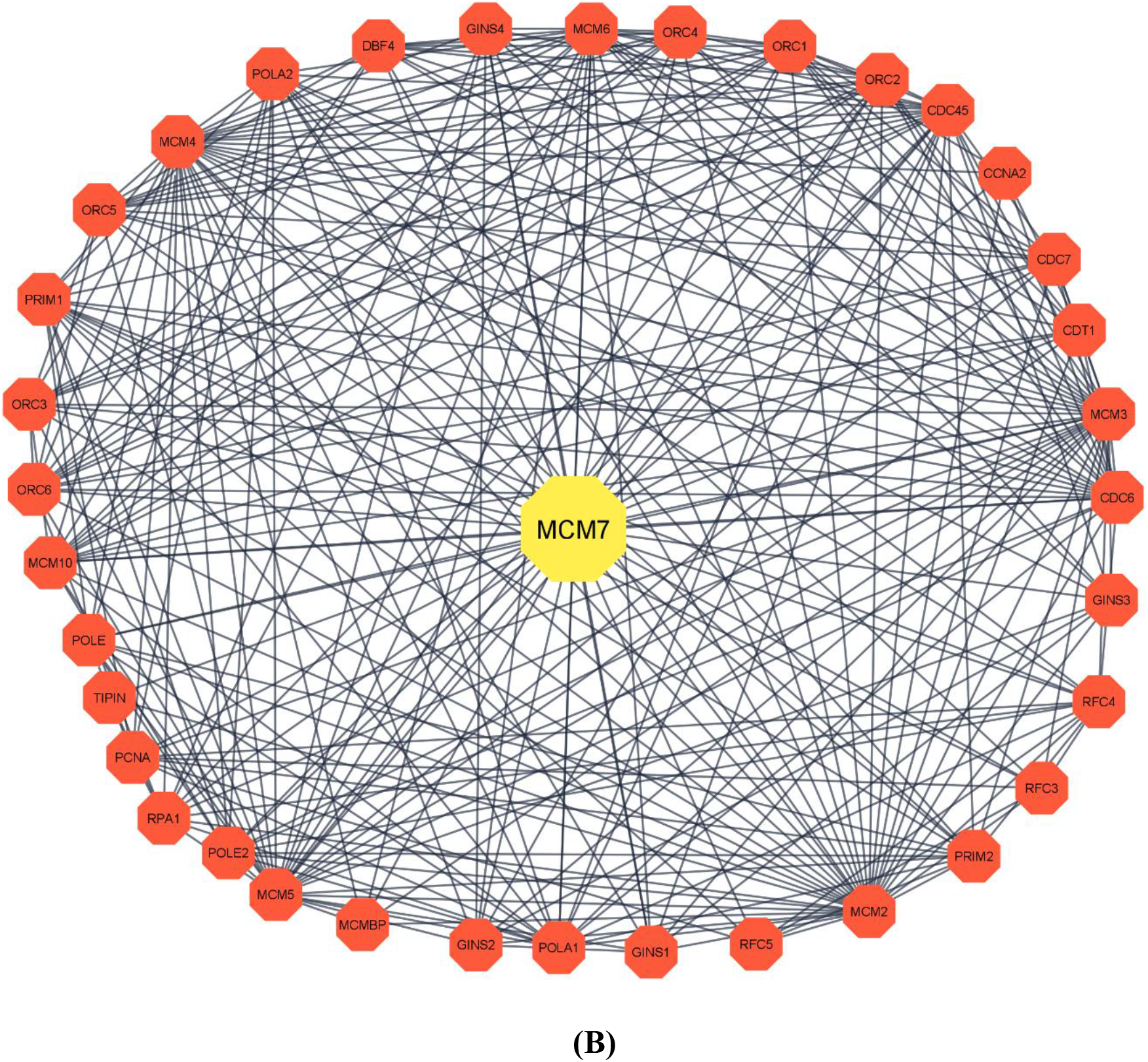
(A) Protein-protein interaction (PPI) network of MCM7 obtained using Cytoscape v3.8.2. (B) PPI Network between MCM7 and its first neighbours (undirected)

### 3.2 Subcellular localization of interacting proteins

Determination of subcellular localization of the target protein and its interacting partners was done using WoLF-PSORT server and a thorough literature survey using UniProt as a source. The results of the analysis have been discussed in the following Table 5 and a Venn diagram, prepared using VENNY 2.1.0 online tool, representing the major intracellular regions of localization of these proteins have been enclosed under ***Figure 5***.

**Table 5:** Predicted subcellular localization of MCM7 and all of its protein-protein interaction partners

| **Protein Name** | **Predicted Subcellular localization** |
| --- | --- |
| **PCNA, DBF4, POLE2, POLE, POLA2, POLA1, PRIM2, PRIM1, CDC7, CCNA1, GINS3, GINS2, GINS1, CDT1, RFC5, RFC4, RFC3, RPA1, MCM7, RPA2, RPA3, MCM8, MCM4, MCM6, MCM2, MCM3, MCM10, MCMBP, ORC1, ORC2, ORC3, ORC4, ORC5, ORC6, CCNE1, LRWD1, E2F1, TOPBP1, CCNL1** | Nucleus |
| **TIPIN, CDC45, CDC6, GINS4, MCM5, GMNN, CHEK1, CCNA2** | Nucleus, Cytoplasm |
| **CDK2, CDK1** | Nucleus, Cytoplasm, Endosomes, Cytoskeleton |
| **MAPRE2** | Cytoskeleton |
| **FEN1** | Nucleus, Mitochondria |

**Figure 5:**
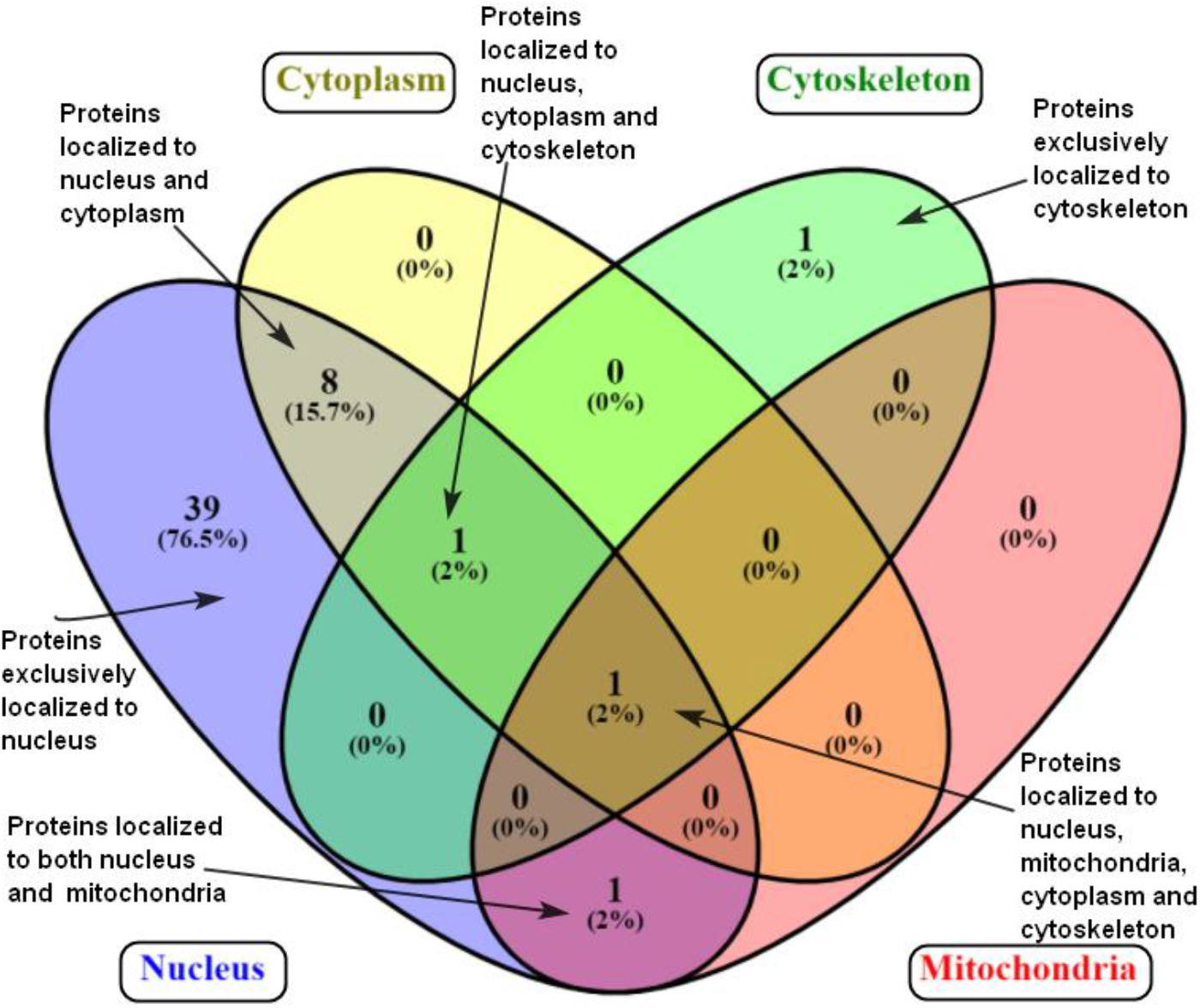
Total percentage of proteins localised in various subcellular compartments-represented by figure generated using Venny 2.1.0

### 3.3. In-silico domain analysis and structure prediction of target protein

For a complete in-silico proteomic analysis of the target protein MCM7, the first step involved the extraction of FASTA sequence of MCM7 from UniProt [22]. Then this FASTA sequence was used as a query to search for its known domains, conserved sites, motifs etc using the EBI INTERPRO [23] server. From the data obtained through UniProt, the major domains and their positions in the protein sequence are as follows: MCM_N (MCM-N terminal domain) (10-139), MCM_OB (oligonucleotide/oligosaccharide binding fold) (149-278), MCM_dom (322-541), MCM_2 (332-537), AAA+_ATPase (ATPases associated with diverse cellular activities), (373-526), AAA_5 (373-526), MCM_lid (558-640) [55]. The secondary structure of the protein was analyzed by using PSIPRED [24] from FASTA sequence to observe properties such as α-helix, β-sheet, turns and coils. The results are represented in Figure(6A) and Figure (6B). Results from additional analysis of data obtained through PSIPRED regarding the reactive nature of the constituent amino acids and their specific properties have been included in the Figure S1(A-C) of supplementary datasheet.

**Figure 6A:**
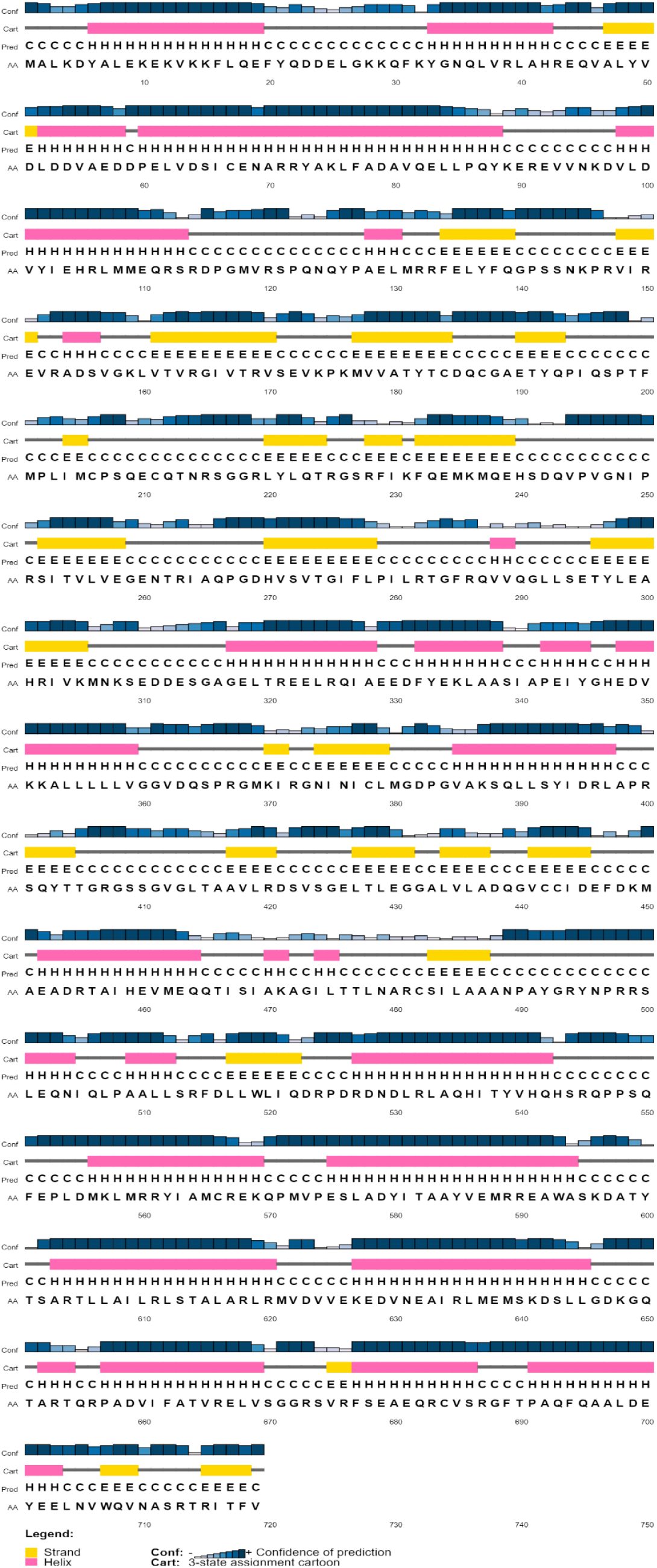
Secondary structure prediction of MCM7 protein

**Figure 6B:**
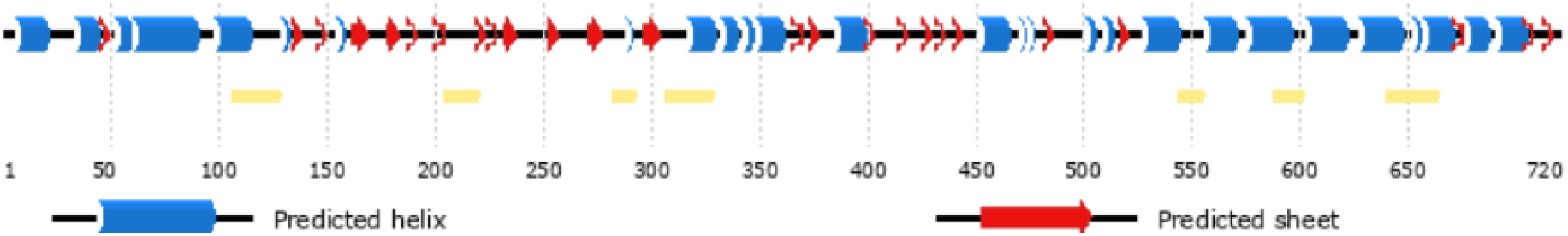
Position dependent feature predictions are mapped onto the sequence schematic shown below. The line height of the Phosphorylation and Glycosylation features reflects the confidence of the residue prediction.

### 3.4 Protein modelling and validation of predicted structures

For performing the homology modelling, the FASTA sequence of then MCM7 protein was computed in BLASTp by using the Protein Data Bank (PDB) database as the search set. The results obtained through BLAST showed only protein structure (present in humans) with sequence similarity above the 80% threshold required for homology modelling. The protein also showed 100% identity with the input FASTA sequence of our target MCM7 protein-therefore ensuring a high degree of accuracy during modelling procedures. The structure for the protein (PDB ID: 6XTX) was extracted from the PDB database. Homology modelling of MCM7 was done using the automated homology protein modelling server of SWISS-MODEL. The user defined template algorithm was selected for computation and the structure extracted from PDB (PDB ID: 6XTX) was used as a template to the target sequence which is MCM7 protein sequence in FASTA format. The MCM7 protein model was verified using the Ramachandran plot from the MolProbity program and validated because the modelled protein’s amino acid residues all fit within the Ramachandran plot’s allowed regions (Figure S2). The MCM7 protein showed 1.17% MolProbity score, 92.33% residues were in the favoured residues, 0.78% in the outliers regions; and the Clash score was 0.59%. On a global scale, the QMEAN Z-score provided an estimate of the “degree of nativeness” of the structural features observed in the model. It indicates whether the model’s QMEAN score is comparable to the expected score from experimental structures of comparable size. QMEAN Z-score value of approximately zero specifies superior quality between the modelled structure and experimental structures. If − 4.0 or lower scores are obtained, then it indicates that the models is of low quality. The QMEAN Z-scores of the MCM7 protein showed − 2.46, and these results indicate that the proposed homology model is reliable and acceptable. The predicted homolog model has been represented in Figure (7A) and Figure (7B). The representations of the analysis of other properties of the modelled protein has been included in the Figure S3 (A-D) of the supplementary data sheet [56–58].

**Figure 7:**
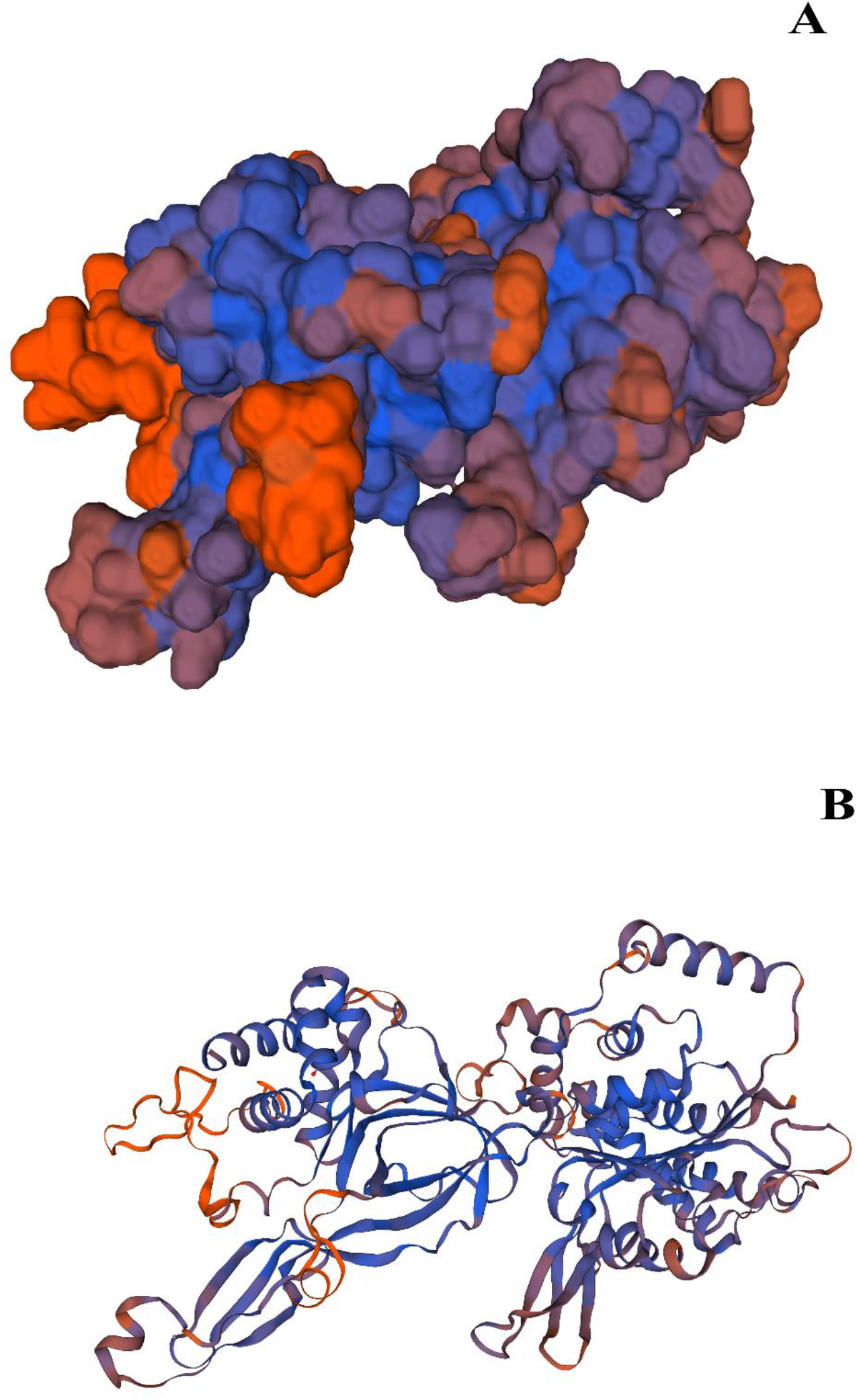
3D representation of predicted MCM7 protein model based on QMEAN parameter generated using SWISSMODEL. **Figure 7(A)** represents the molecular surface structure and **Figure 7(B)** represents the thread-ribbon structure. The colour grading is as follows: Orange= low confidence prediction/low QMEAN score/low sequence similarity with template structure, Purple= moderate confidence prediction/moderate QMEAN score/moderate sequence similarity with template structure, Blue= high confidence prediction/high QMEAN score/high sequence similarity with template structure.

The results obtained through SWISSMODEL were validated using SAVESv6.0 server through PROCHECK and VERIFY3D algorithms. The results obtained were satisfactory to validate the predicted model. The results are depicted in Figure (8A) and Figure (8B).

**Figure 8A:**
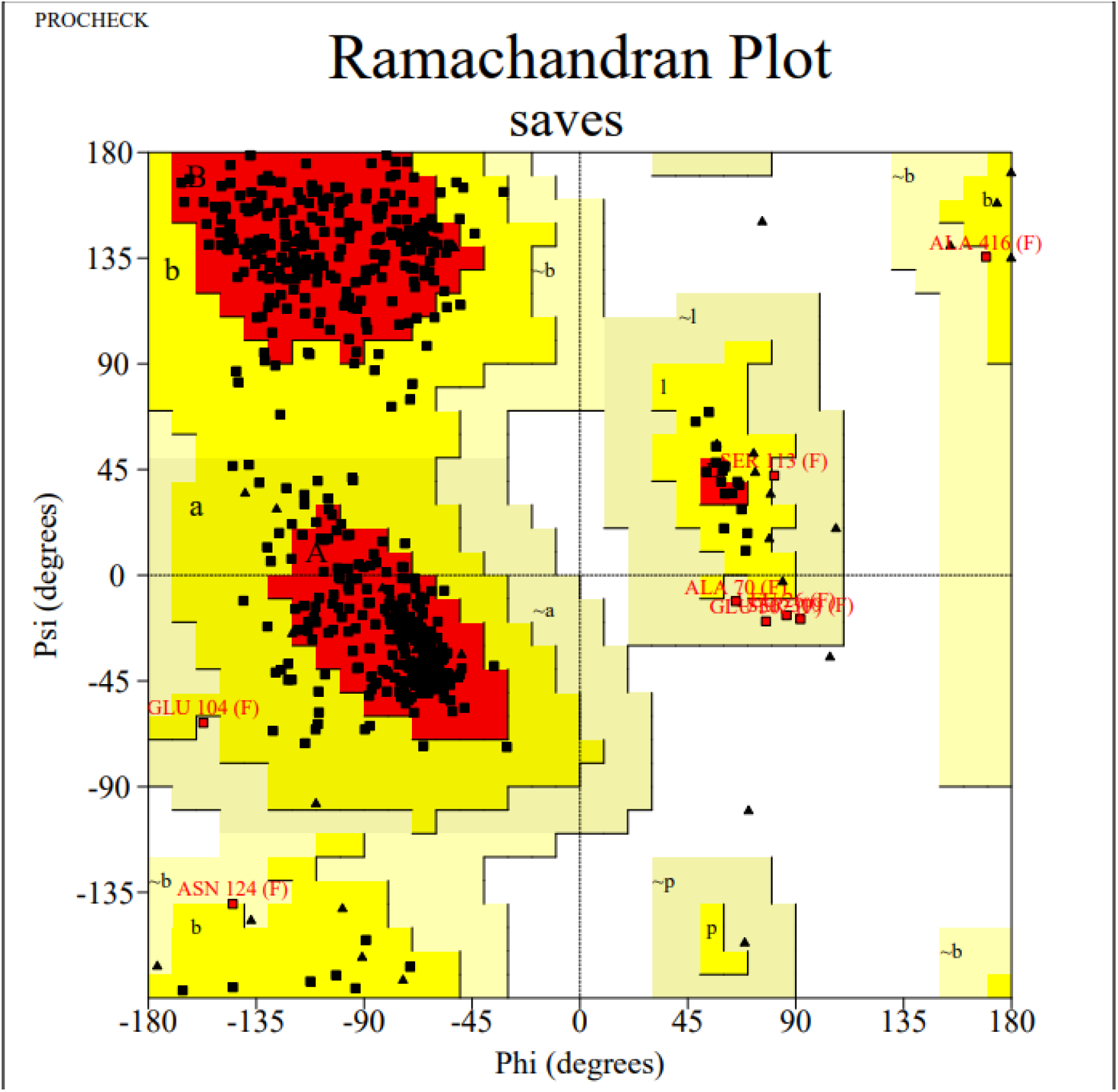
Validation of predicted structure of MCM7 protein using PROCHECK

**Figure 8B:**
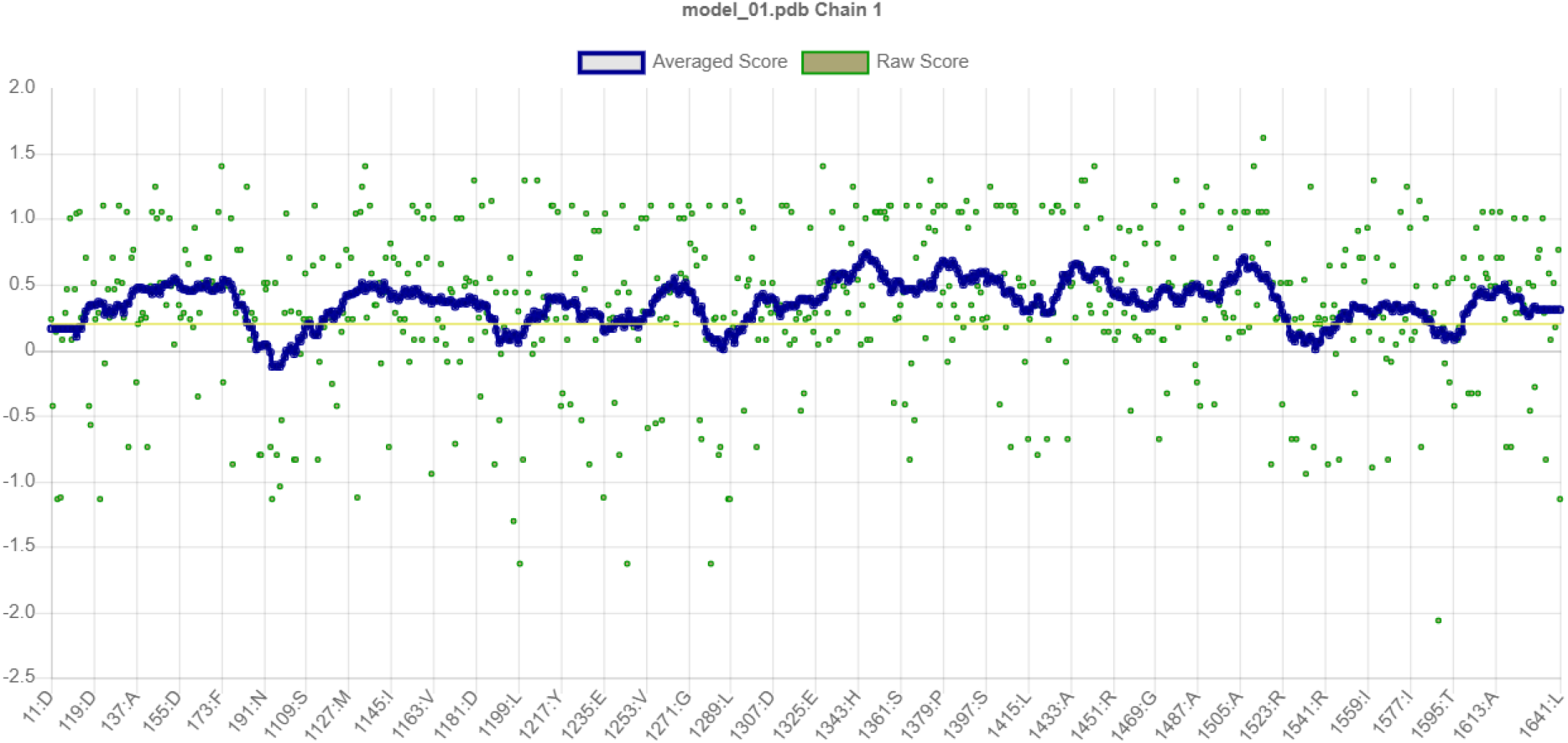
Validation of predicted structure of MCM7 protein using VERIFY3D

For the purpose of performing further validation, the obtained predicted structure was compared against the structure for the MCM7 protein obtained through PDB using the PROMALS3D online server. Although the multiple sequence alignment results in PROMALS3D show the presence of a few minor gaps in the alignment of the two sequences, phylogenetic guide tree results ***(Figure 9)*** show that these two protein structures are represented on equidistant nodes from the root node, confirming the high degree of similarity or identity between the two sequences.

**Figure 9:**
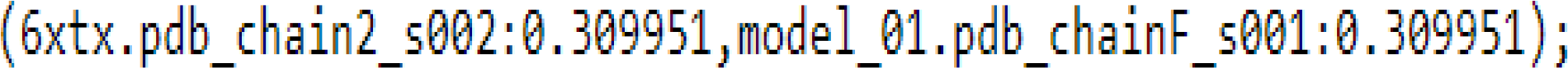
PROMALS3D phylogenetic tree prediction (6XTX represents PDB model template; model_01 represents structure predicted by SWISS-MODEL

## 4. CONCLUSION AND FUTURE PROSPECTS

MCM7 is upregulated in Alzheimer’s Disease and downregulated in the three grades of Breast Cancer as per our analysis. This gene forms a key component in the DNA replication process and cell cycle progression. Thus, any mutation in this gene or any structural changes in the subsequent protein affects the DNA replication process as well as the cell cycle, thereby increasing the probability of occurrence of cellular aberrations such as onset of cancer or changes in protein structure or folding resulting in onset of diseases such as Alzheimer’s. In order to assess therapeutic options for breast cancer and/or Alzheimer’s Disease in a shorter time-scale without using much capital investment on R&D of novel drugs and toxicological research, drug repurposing studies can be done in order to observe the effect of pre-existing drugs on rectifying expression levels of a mutated MCM7 gene. Based on the above-mentioned findings, novel drugs can also be synthesized which can act on rectifying any erroneous expression of the MCM7 gene.

## ACKNOWLEDGEMENT

The authors acknowledge the COVID-19 pandemic to motivate for switching research work to computational biology and further extend gratitude towards the MHRD (Govt. of India) for financial support to the first author.

